# *In silico* analysis predicts a limited impact of SARS-CoV-2 variants on CD8 T cell recognition

**DOI:** 10.1101/2022.03.23.485487

**Authors:** Olga I. Isaeva, Steven L.C. Ketelaars, Pia Kvistborg

## Abstract

Since the start of the COVID-19 pandemic, mutations have led to the emergence of new SARS-CoV-2 variants, and some of these have become prominent or dominant variants of concern. This natural course of development can have an impact on how protective the previously naturally or vaccine induced immunity is. Therefore, it is crucial to understand whether and how variant specific mutations influence host immunity. To address this, we have investigated how mutations in the recent SARS-CoV-2 variants of interest and concern influence epitope sequence similarity, predicted binding affinity to HLA, and immunogenicity of previously reported SARS-CoV-2 CD8 T cell epitopes. Our data suggests that the vast majority of SARS-CoV-2 CD8 T cell recognized epitopes are not altered by variant specific mutations. Interestingly, for the CD8 T cell epitopes that are altered due to variant specific mutations, our analyses show there is a high degree of sequence similarity between mutated and reference SARS-CoV-2 CD8 T cell epitopes. However, mutated epitopes, primarily derived from the spike protein, in SARS-CoV-2 variants Delta, AY.4.2 and Mu display reduced predicted binding affinity to their restriction element. These findings indicate that the recent SARS-CoV-2 variants of interest and concern have limited ability to escape memory CD8 T cell responses raised by vaccination or prior infection with SARS-CoV-2 early in the pandemic. The overall low impact of the mutations on CD8 T cell cross-recognition is in accordance with the notion that mutations in SARS-CoV-2 are primarily the result of receptor binding affinity and antibody selection pressures exerted on the spike protein, unrelated to T cell immunity.

## 1. Introduction

The COVID-19 pandemic caused by SARS-CoV-2 is having a global catastrophic impact on public health and social economy (1,2). SARS-CoV-2 was first identified in humans in late 2019 in Wuhan, China, and the outbreak was designated as a pandemic by the WHO on March 11th, 2020 (3,4). The early variant of SARS-CoV-2 (also known as lineage B or Wuhan-Hu-1; UniProt: UP000464024; Genome accession: MN908947) is hereafter referred to as ‘reference SARS-CoV-2’.

SARS-CoV-2 is a single-stranded RNA virus characterized by an inherently high mutation rate, short replication time and high virion yield (5–8). As the virus spreads, this leads to a high genetic diversity and allows the virus to evolve rapidly as a result of natural selection pressures, including those originating from the host immune system. Mutations accumulate over time and result in amino acid changes that decrease the antigenicity of immune targeted proteins. This gradual change in antigenicity of viral proteins, driven by selective immune pressure, is known as antigenic drift (9). Antigenic drift allows viruses to continuously evade host immunity, facilitating recurrent viral outbreaks. In acute infectious disease, antigenic drift is primarily driven by antibody responses leading to selection of escape mutants (9). In accordance with this, many of the amino acid changes in SARS-CoV-2 variants are located in the spike protein, the main target of neutralizing antibodies (10). These antibodies form the only immune mechanism that is able to provide sterilizing immunity, preventing host cells from being infected. The rate of evolution of the SARS-CoV-2 spike protein is much higher than that of similar proteins in other known viruses that cause acute infectious disease in humans. For example, its rate of evolution is approximately 10-fold higher than the evolution rate of the influenza A hemagglutinin and neuraminidase proteins (9). In addition, a large number of amino acid changes have accumulated in SARS-CoV-2 proteins that are not known antibody targets (11). These amino acid changes may have inferred a fitness advantage to the virus unrelated to antibody immunity, as antigenic drift is primarily driven by antibody responses in acute viral infections (6,9,12,13).

Even though T cells are unlikely to be a main source to antigenic drift there is ample evidence for the importance of these cells in protection against severe and critical COVID-19 and re-infections: 1) Depletion of CD8 T cells led to impaired clearance of SARS-CoV-2 in a COVID-19 mouse model(14) and breakthrough infections in a rhesus macaque model upon rechallenge; 2) Lower baseline peripheral blood CD8 T cell counts have been shown to correlate with decreased patient survival (15,16); and 3) CD8 T cells have also been shown to impact COVID-19 disease severity: high percentages of HLA-DR+CD38hi CD8+ T cells in peripheral blood of COVID-19 patients were demonstrated to correlate with disease severity (17), and early bystander CD8 T cell activation combined with absence of systemic inflammation was shown to predict asymptomatic or mild disease (18). Combined, these observations suggest that CD8 T cell immunity is important for protection against reinfection and severe COVID-19 disease (19).

As a direct consequence of antigenic drift, several SARS-CoV-2 variants defined by amino acid changes that directly impact virus transmissibility, pathogenicity, infectivity and/or antigenicity have emerged (20). The most prominent SARS-CoV-2 variants in Europe were designated as variants of concern (VOC) (Alpha, B.1.1.7; Beta, B.1.351; Gamma, P.1; Delta, B.1.617.2; Omicron, B.1.1.529) and variants of interest (VOI) (Lambda, C.37; Mu, B.1.621; “Delta Plus”, AY.4.2) according to the European Centre for Disease Prevention and Control (ECDC) designation (21). All VOC and VOI except AY.4.2 were also designated as VOC or VOI by the World Health Organization at the moment of this investigation (20). SARS-CoV-2 variant Alpha was the dominant variant in circulation starting in late 2020 and was subsequently replaced by SARS-CoV-2 variant Delta which accounted for 90% of the infections worldwide by August 2021. In November 2021, SARS-CoV-2 variant Omicron was first detected. It was responsible for at least 92% of global SARS-CoV-2 infections by February 2022 **(Figure 1A**, (22)). There is accumulating evidence that recent SARS-CoV-2 variants including Beta, Delta and Omicron are less efficiently neutralized by vaccine recipients’ sera (23,24). In terms of T cell immunity, there is experimental data by other groups showing that T cell responses induced by reference SARS-CoV-2 generally cross-recognize SARS-CoV-2 variants Alpha, Beta, Gamma, Delta and Omicron (25–29). However, these papers do not include systematic data regarding the effect of SARS-CoV-2 variant-specific amino acid changes on the properties of previously recognized CD8 T cell epitopes.

**Figure 1.**
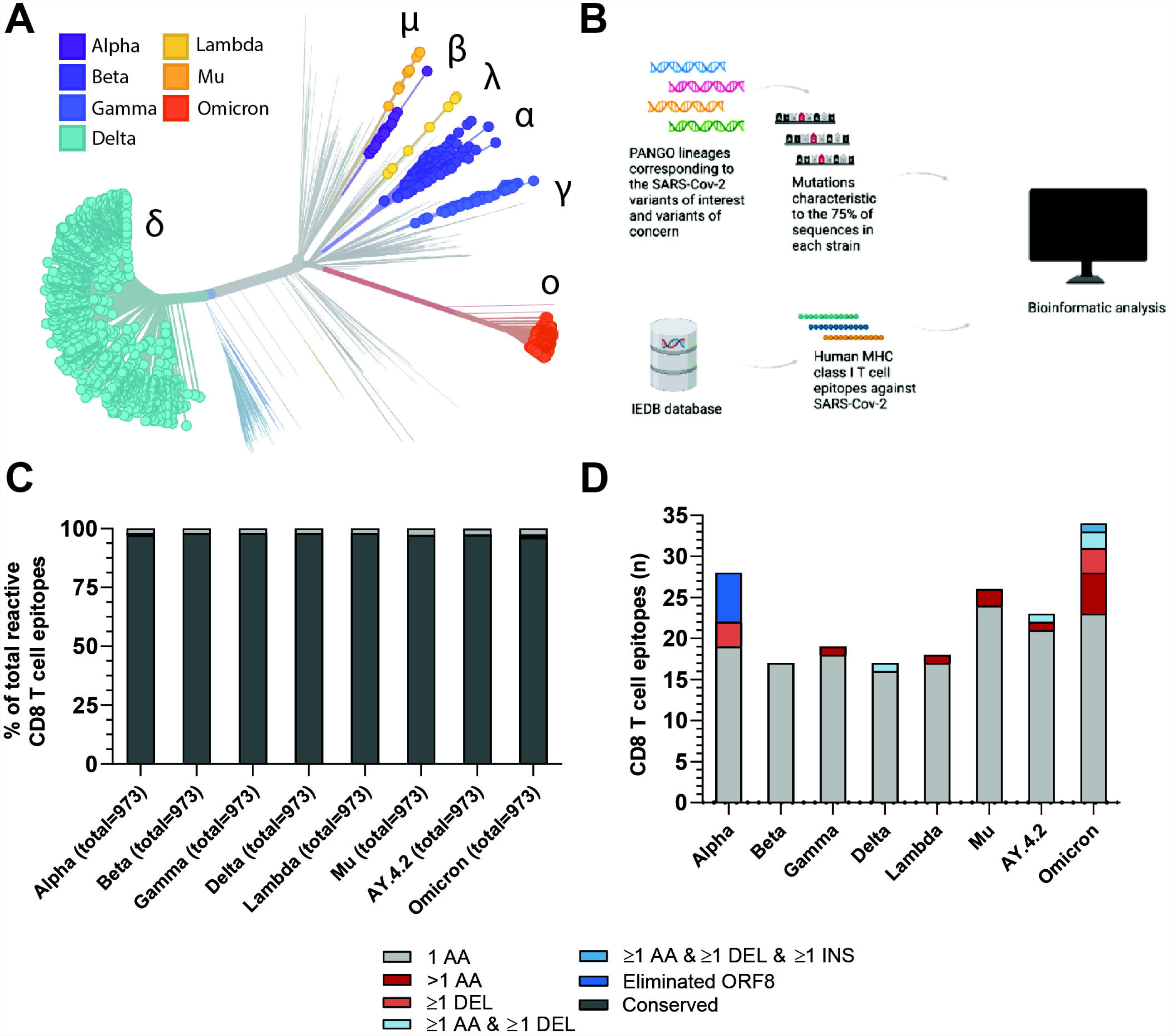
Overview of the investigated SARS-CoV-2 CD8 T cell reactive epitopes. **(A)** Phylogenetic tree where isolates originating from variants of concern (VOCs) Alpha, Beta, Gamma, Delta and Omicron are highlighted, as well as variants of interest (VOIs) Lambda and Mu. AY.4.2 is a subvariant of Delta and overlaps with the Delta branch. The length of the branches reflects the time of emergence. Visualization generated using the Nextstrain platform (22). **(B)** Depiction of the methods used in the project. **(C)** Stacked bar graph indicating the percentages of CD8 T cell recognized epitopes per variant that are conserved or harbor the indicated types of mutations. **(D)** Numbers of CD8 T cell recognized epitopes per variant that harbor the indicated types of mutations.

In this work, we investigate the potential consequences of variant specific mutations on the SARS-CoV-2 specific CD8 T cell responses raised by either natural infection or vaccination based on in silico analysis. In particular, we explore changes in predicted binding affinity of the epitopes to their HLA restriction elements, predicted immunogenicity and likelihood of CD8 T cell receptor cross-recognition of epitopes between the reference SARS-CoV-2 strain and SARS-CoV-2 variants of interest and concern. We perform these analyses pan-proteome to identify the degree of protection after a natural infection. Furthermore, the vaccines currently approved by WHO are limited to the spike protein (30). Therefore, we have also conducted the analyses focused on CD8 T cell recognized epitopes derived from the spike protein only to determine the degree of the vaccine-mediated protection.

## 2 Materials and methods

### 2.1 Identification of dominant non-synonymous mutations in SARS-CoV-2 variants

The list of SARS-CoV-2 variants of interest and variants of concern has been compiled according to the WHO and ECDC designations as of December 10, 2021. For each of the variants, a list of mutations present in 75% of the GISAID sequences for the corresponding PANGO lineage was compiled via the outbreak.info API. The lists of mutations per lineage can be found in Supplementary Table 1 (31,32).

### 2.2 Parsing of CD8 T cell recognized SARS-CoV-2 epitopes using IEDB

The table of T cell assay results was downloaded from the IEDB website on December 8, 2021 (33,34). The table was filtered to include only linear SARS-CoV-2 epitopes in humans, presented in context of the MHC class I. Only epitopes from patients with infectious disease were included. Only positive assays with negative adoptive flag field were included. The tables of variant mutations and SARS-CoV-2 CD8 T cell recognized epitopes were subsequently intersected. CD8 T cell recognized epitopes that were deduced from reactive overlapping peptide pools were filtered from the list. Epitopes with published HLA restriction elements were manually curated. The final list of epitopes and corresponding HLA alleles is shown in the Supplementary Tables 2 and 3.

### 2.3 Epitope analysis

The normalized epitope similarity score between the altered and reference SARS-CoV-2 CD8 T cell recognized epitope was calculated as described by Frankild et al. (35). This method does not allow the calculation of the similarity score between two sequences of differing lengths. For this reason, we have set the sequence similarity score of CD8 T cell recognized epitopes harboring deletions and/or insertions to 0. IEDB’s epitope cluster analysis tool was additionally used on each reference and altered epitope to determine if the epitope pairs share a sequence identity of 80%, 80-90% or more than 90% (36). The parameters used were: minimum sequence identity threshold: 80%, 90%. Minimum/Maximum peptide length: NA. Clustering method: fully interconnected clusters (cliques).

IEDB’s T cell class I pMHC immunogenicity prediction tool was used to compare the immunogenicity of the altered and reference SARS-CoV-2 CD8 T cell recognized epitopes (37). The default setting was used, masking the 1st, 2nd and C-terminus amino acids of the epitopes in the analysis.

For all parsed reference SARS-CoV-2 CD8 T cell recognized epitopes with experimentally validated HLA restriction information, the predicted binding affinity to the given HLA was calculated for both the reference and altered epitope using NetMHCpan-4.1 (38). The predicted binding affinity is expressed as the half-maximal inhibitory concentration IC50 nM. For each paired reference and altered epitope, the fold change in predicted binding affinity as a result of the mutation(s) was calculated. A 2-fold change in predicted binding affinity was defined as a decrease in predicted binding affinity, a fold change below 0.5 as an increase in predicted binding affinity and a fold change between 0.5 and 2 was conservatively defined as neutral. CD8 T cell recognized epitopes overlapping with a deletion and/or insertion and not predicted to bind to the HLA as a result of the mutation were defined as decreased in predicted binding affinity.

### 2.4 Statistical analysis

For all analyzed SARS-CoV-2 CD8 T cell recognized reference and altered epitope pairs, differences in predicted immunogenicity and predicted binding affinity were assessed using a two-tailed Wilcoxon matched-pairs signed rank test. The increase in fractions of CD8 T cell recognized epitopes with decreased binding affinity and/or an epitope sequence similarity <85% was also assessed using a two-tailed Wilcoxon matched-pairs signed rank test. Comparisons in log2 fold change predicted binding affinity and/or epitope sequence similarity between spike and non-spike protein-derived mutated CD8 T cell recognized epitopes were assessed using a two-tailed Mann–Whitney *U* test. Differences were considered significant if *P*□<□0.05. Statistical analysis was performed with GraphPad Prism (version: 8.4.2, for Windows, GraphPad Software, San Diego, California USA, (39)).

## 3 Results

### 3.1 A minor fraction of SARS-CoV-2 derived CD8 T cell recognized epitopes are mutated in Variants of Concern and Interest

We focused our analysis on the current SARS-CoV-2 variants of concern (VOC) (Alpha, B.1.1.7; Beta, B.1.351; Gamma, P.1; Delta, B.1.617.2; Omicron, B.1.1.529) and variants of interest (VOI) (Lambda, C.37; Mu, B.1.621; “Delta Plus”, AY.4.2). First, we identified the non-synonymous amino acid substitutions, insertions and deletions that were present in at least 75% of total virus isolates for each variant in the GISAID database (per December 6^th^, 2021; (31,40), as compared to the reference SARS-CoV-2 variant (Supplementary Table 1). Next, we parsed all 973 unique experimentally validated CD8 T cell recognized reference SARS-CoV-2 derived epitopes identified in patients with COVID-19, per December 8^th^ 2021, from the Immune Epitope Database (IEDB) (34,41) and aligned these with the sequences spanning the identified non-synonymous mutations in the investigated SARS-CoV-2 variants (Figure 1B). Specifically, all SARS-CoV-2 CD8 T cell epitopes detected in patients with COVID-19 were included. Epitopes deduced from peptide pools, in which the exact reactive peptide is not validated, were filtered out. In addition, studies conducted in the adoptive transfer setting were filtered out.

The vast majority of the 973 included CD8 T cell recognized epitopes was found to be conserved across the different variants (median: 97.8%, range: 96.5-98.3%): we identified a total of 93 unique epitopes that harbored one or more mutations (Figure 1C). Specifically, between 17 and 34 unique epitopes per variant (median: 21) overlap with one or more amino acid substitutions, deletions and/or insertions (Figure 1D, Supplementary Tables 2 and 3). Six CD8 T cell recognized epitopes were considered eliminated in SARS-CoV-2 variant Alpha as they were located downstream of a stop-codon mutation (ORF8 Q27*); three additional epitopes contain a deletion (S∆69/70 or ∆144/144). In SARS-CoV-2 variant Beta, the identified CD8 T cell recognized epitopes only contain single amino acid substitutions. Altered CD8 T cell recognized epitopes in the more recent Gamma, Delta, Lambda, Mu and AY.4.2 SARS-CoV-2 variants do not harbor single deletions but harbor other types of mutations, for example, epitopes with mutations consisting of more than one amino acid substitution (Gamma, Lambda, Mu and AY.4.2; n = 1, 1, 2 and 1, respectively) or an epitope with a deletion (S∆157-158) together with an amino acid substitution (Delta and AY.4.2). The recently emerged SARS-CoV-2 Omicron variant harbors the largest number of CD8 T cell recognized epitopes that overlap with non-synonymous mutations (n=34). These mutations result in epitopes with single (n=23), double (n=2) and triple (n=3) amino acid substitutions; single deletions (n=3); a combined amino acid substitution and deletion (n=2), and even a combined substitution, deletion and insertion (n=1) (Figure 1D, Supplementary Tables 2 and 3).

To be able to investigate the potential consequences of the variant specific mutations on T cell recognition, we made a list of all variant specific CD8 T cell epitopes based on the variant specific mutations. For the analyses investigating the likelihood of T cell receptor cross-recognition and epitope immunogenicity of the altered epitopes, we included the 93 unique CD8 T cell recognized epitopes with variant specific mutations. For the prediction of HLA binding affinity, we limited the analysis to the 74 of the 93 epitopes for which HLA restriction elements had been experimentally determined by the scientific community (Supplementary Tables 2 and 3). In total, these epitopes bind 27 HLA alleles, with between 1 and 14 epitopes per allele (median: 3, Figure S1A).

### 3.2 Properties of altered epitopes are highly conserved between variants and reference SARS-CoV-2 CD8 T cell recognized epitopes

Amino acid changes in SARS-CoV-2-derived CD8 T cell epitopes can reduce the sequence similarity to the reference epitope. The more distinct the biochemical properties of an amino acid substitution are compared to the reference amino acid, the greater the dissimilarity. This could lead to reduced or abrogated activation of memory CD8 T cells reacting to the altered epitope. The epitope sequence similarity of the altered epitope to the reference epitope can therefore be used as an *in silico* proxy for likelihood of T cell receptor cross-recognition.

To test the epitope sequence similarity between the variant-specific and matched reference epitopes, we conducted two analyses: 1) We compared the sequence similarity between the reference and the altered epitopes using a previously published method (35). Importantly, this method incorporates the biochemical properties of amino acid substitutes to score the epitope sequence similarity, which is crucial in epitope cross-recognition by CD8 T cell receptors. Experimental data demonstrate that CD8 T cell receptor recognition drops to 50% if peptide similarity drops below 85% (42). We found that the vast majority (median: 90%, range: 65-100%) of the reference and the matched variant specific CD8 T cell epitopes share over 85% sequence similarity (Figure 2A and S2A). 2) In addition, we measured the degree of sequence similarity between the pairs of epitopes using the IEDB clustering tool which performs a local alignment (36). In contrast to the first method, the IEDB clustering tool allows comparison of epitopes of differing lengths (e.g. due to insertions/deletions). Data from our previous experiments in the tumor setting suggests that a sequence similarity above 80% could serve as an indicator of potential TCR cross-reactivity (43,44). The majority (median: 93%, range: 73-100%) of reference SARS-CoV-2 epitopes and variant derived epitopes share at least 80% similarity (Figure S2B). Taken together, these *in silico* analyses suggest that the ability of memory CD8 T cells, induced by natural infection with the reference virus, to respond to the included variants is not significantly impaired.

**Figure 2.**
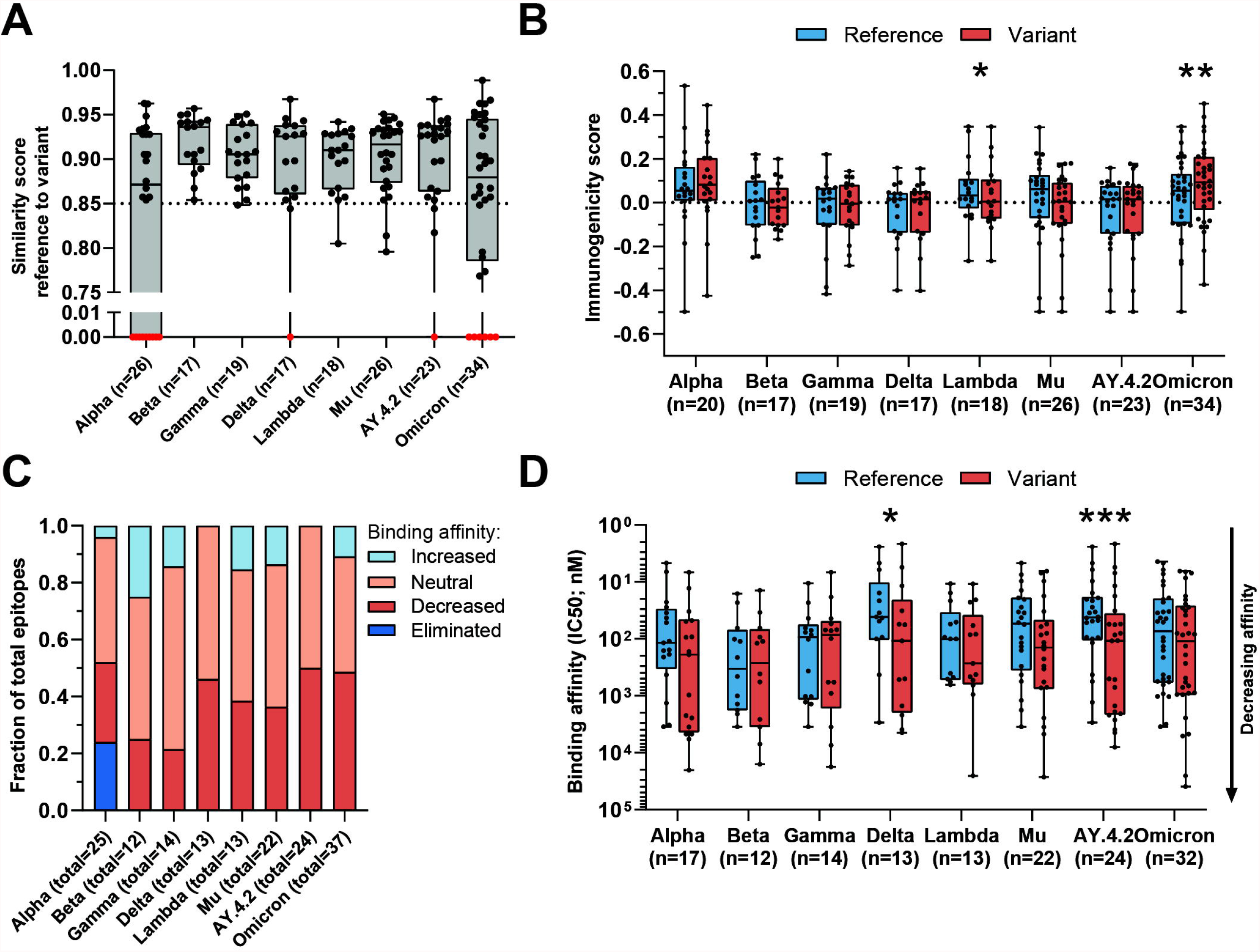
Sequence similarity and predicted binding affinity of mutated CD8 T cell recognized epitopes. **(A)** Sequence similarity scores between the reference SARS-CoV-2 CD8 T cell recognized epitopes and the altered epitopes. Sequence similarity of epitopes in red is set to zero as a result of one or more deletions/insertions in the epitope sequence (Alpha, n = 3; Delta, n = 2; AY.4.2, n = 2; Omicron, n = 13) or due to the ORF8 Q27* stop codon mutation (Alpha, n = 6). **(B)** Box plot indicating the predicted immunogenicity of the reference SARS-CoV-2 CD8 T cell recognized epitopes and the altered epitopes according to the IEDB T cell class I pMHC immunogenicity prediction tool. **(C)** Fractions of total altered CD8 T cell recognized epitopes where the predicted binding affinity of the epitope to the corresponding HLA restriction element was increased (≤0.5-fold change in IC50), remained neutral (0.5< fold change in IC50 <2) or was decreased (≥2-fold change in IC50) as a result of the mutation. Epitopes were considered eliminated as a result of the ORF8 Q27* stop codon mutation (Alpha variant). **(D)** Box plot indicating the predicted binding affinity IC50 (nM) of the reference and altered CD8 T cell recognized epitope to the corresponding HLA restriction element. Box plots indicate the median (line), 25th and 75th percentile (box), min and max (whiskers), and all data points (single circles). Statistical significance was tested with a two-tailed Wilcoxon matched-pairs signed rank test. Variation in numbers of epitopes in the analyses are due to inclusion of epitopes binding one or more HLA restriction elements. * P < 0.05, *** P < 0.001. n indicates the number of epitopes analyzed per group.

The likelihood that a certain peptide is immunogenic can be predicted based on the presence and, importantly, positioning of amino acids with certain biochemical properties (37). To investigate whether a SARS-CoV-2 CD8 T cell recognized epitope is predicted to be more or less immunogenic as a result of an amino acid change, we applied the IEDB T cell class I pMHC immunogenicity prediction tool to the set of reference SARS-CoV-2 CD8 T cell recognized epitopes and variant derived epitopes. This tool uses a large set of known peptide immunogenicity values to computationally predict whether CD8 T cell epitopes are immunogenic (i.e., likelihood for T cell recognition) or not (37). Surprisingly, the epitopes derived from the Omicron and Lambda variants were predicted to be significantly more and less immunogenic, respectively (Omicron: p=0.0042, (Lambda: p=0.03; Figure 2B). For all included SARS-CoV-2 variants, a large fraction of mutated epitopes was predicted to be either more immunogenic (median: 47%, range: 11-57%) or unchanged in immunogenicity (median: 28%, range: 19-44%). Between 6% and 44% (median: 24%) of variant specific CD8 T cell recognized epitopes were predicted to be less immunogenic as a result of the mutation (Figure S3A). Taken together, these analyses indicate that there is a high degree of sequence similarity between altered and reference epitopes in all analyzed SARS-CoV-2 variants, which is likely to result in a high degree of CD8 T cell cross-reactivity between these epitopes.

### 3.3 A minor fraction of mutated epitopes from Delta and AY.4.2 exhibit reduced predicted binding affinity to MHC class I

Amino acid changes in CD8 T cell recognized epitopes may result in altered binding affinity to the corresponding HLA restriction elements. This may result in altered presentation of the epitope on the surface of SARS-CoV-2 infected cells, making the infected cells less visible to T cell recognition. To estimate the changes in binding affinity of the altered epitopes, we used the 74 unique SARS-CoV-2 CD8 T cell recognized epitopes with previously experimentally validated HLA restriction elements. We used the NetMHCpan-4.1 tool to predict the binding affinity of each reference and variant specific CD8 T cell epitope to the matched HLA restriction element (38). In this analysis, epitopes that can bind to more than one HLA allele were included for each of the HLA allele they bind to.

For each included SARS-CoV-2 variant, we observed decreased binding affinity, defined as a ≥2-fold change in IC50 value for a subset of the variant specific epitopes (median 37% of epitopes, range: 21%-50%) (Figure 2C). Between 41% and 64% (median: 50%) of variant specific epitopes retained their predicted binding affinity (neutral; 0.5< fold change in IC50 <2), and for between 0% and 25% (median: 12%) of altered epitopes an increased binding affinity was predicted (≤0.5-fold change in IC50 value). Following a comparison of the difference in predicted binding affinity of the paired reference SARS-CoV-2 CD8 T cell recognized epitopes and mutated epitopes, the small set of epitopes of the Delta variant and its subvariant AY.4.2 were predicted to have a significantly reduced binding affinity to the HLA as a result of their mutations (Delta: p=0.01, AY.4.2: p=0.0002; Figure 2D). Importantly, despite these statistically significant differences, these results are derived from a highly limited number of epitopes (12 and 18 altered epitopes derived from Delta and AY.4.2, respectively, out of a total of 973 CD8 T cell recognized epitopes per variant).

### 3.4 A larger fraction of spike derived CD8 T cell epitopes are affected by variant specific mutations compared to pan proteome derived epitopes

To date, it is estimated that since the start of the pandemic there have been more than 400 million COVID-19 cases (45). This translates to approximately 5% of the world population, however, many cases were never included in the official statistics. In contrast, it is estimated that over half of the world population (62.6% on February 25, 2022) has received at least one dose of a COVID-19 vaccine based on the reference SARS-CoV-2 sequence of the spike protein (46). Of all the proteins encoded by SARS-CoV-2, the spike protein is subject to the highest rate of evolution (10). As a consequence, spike protein-derived CD8 T cell recognized epitopes are inherently the least conserved T cell epitopes of the SARS-CoV-2 proteome. Individuals that have not been infected but have only received the vaccine may have a lower level of protective CD8 T cell immunity due to loss of the part of the epitopes.

We performed our analysis on spike protein-derived epitopes only. The SARS-CoV-2 spike protein encodes 263 previously identified CD8 T cell recognized epitopes. The majority of these 263 CD8 T cell recognized epitopes is conserved across the variants included in our analysis (median: 95.1%, range: 92.0-96.6%) corresponding to between 9 and 21 (median: 13) epitopes per variant which have alterations in the amino acid sequence (Figure 3A, Figure S1C, Supplementary Tables 2 and 3). The majority (median 84%, range: 52-100%) of these mutated variant specific epitopes share at least 85% similarity with the corresponding references epitopes (Figure 3B and S2C) based on the IEDB epitope clustering tool (median 89%, range: 71-100%; figure S2D).

**Figure 3:**
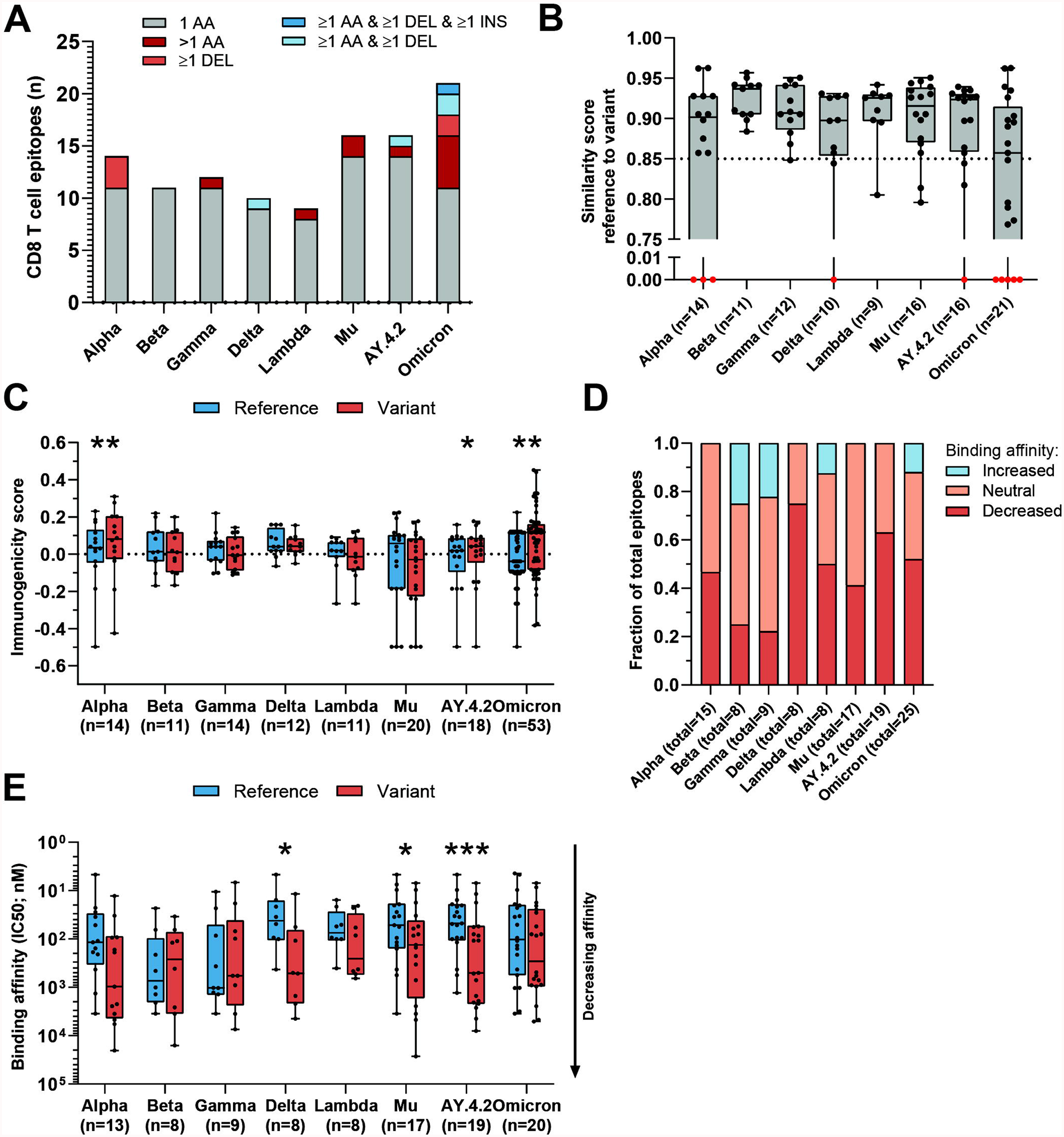
Sequence similarity and predicted binding affinity of mutated CD8 T cell recognized epitopes derived from the spike protein. **(A)** Numbers of spike protein-derived CD8 T cell recognized epitopes per variant that harbor the indicated categories of mutations. **(B)** Sequence similarity scores between the spike protein-derived reference SARS-CoV-2 CD8 T cell recognized epitopes and the matched variant epitopes. Sequence similarity of epitopes in red is set to zero as a result of one or more deletions/insertions in the epitope sequence (Alpha, n = 3; Delta, n = 2; AY.4.2, n = 2; Omicron, n = 12) **(C)** Box plot indicating the predicted immunogenicity of the spike protein-derived reference SARS-CoV-2 CD8 T cell recognized epitopes and the altered epitopes according to the IEDB T cell class I pMHC immunogenicity prediction tool. **(D)** Fractions of total altered spike protein-derived CD8 T cell recognized epitopes where the predicted binding affinity of the epitope to the corresponding HLA restriction element was increased (≤0.5-fold change in IC50), remained neutral (0.5< fold change in IC50 <2) or was decreased (≥2-fold change in IC50) as a result of the mutation. **(E)** Box plot indicating the predicted binding affinity IC50 (nM) of the reference and altered spike protein-derived CD8 T cell recognized epitope to the corresponding HLA restriction element. Box plots indicate the median (line), 25th and 75th percentile (box), min and max (whiskers), and all data points (single circles). Statistical significance was tested with a two-tailed Wilcoxon matched-pairs signed rank test. Variation in numbers of epitopes in the analyses are due to inclusion of epitopes binding one or more HLA restriction elements. AA: amino acid, DEL: deletion, INS: insertion. * P < 0.05, ** P < 0.01, *** P < 0.001. n indicates the number of epitopes analyzed per group.

Interestingly, the mutated spike protein-derived epitopes from the Alpha, AY.4.2 and Omicron variants are predicted to be significantly more immunogenic compared to reference (Alpha: p=0.0034, AY.4.2: p=0.031, Omicron: p=0.0065; Figure 3C). A large fraction of mutated epitopes was predicted to be either more immunogenic (median: 63%, range: 22-79%) or unchanged in immunogenicity (median: 20%, range: 11-31%). Between 6% and 67% (median: 15%) of variant specific CD8 T cell recognized epitopes were predicted to be less immunogenic as a result of the mutation (Figure S3B). Furthermore, for each included SARS-CoV-2 variant, decreased binding affinity is predicted (≥2-fold change in IC50 value; Figure 3D) for a subset of the altered epitopes (median 48% of epitopes, range: 22%-75%). Between 25% and 59% (median: 44%) of altered epitopes retained their predicted binding affinity (neutral; 0.5< fold change in IC50 <2), and between 0% and 25% (median: 6%) of altered epitopes had an increased predicted binding affinity (≤0.5-fold change in IC50 value). Spike protein-derived CD8 T cell recognized epitopes of the Delta, Mu and AY.4.2 variants were predicted to have a significantly reduced binding affinity to the HLA as a result of their mutations (Delta: p=0.016, Mu: p=0.017, AY.4.2: p=0.0002; Figure 3E). Importantly, despite these statistically significant differences, these results are derived from only 8 to 14 (median: 12) unique CD8 T cell recognized epitopes that are mutated per variant, out of a total of 263 unique epitopes per variant.

Next, we tested whether variant-specific mutations may have a more profound effect on spike protein-derived CD8 T cell recognized epitopes compared to non-spike. We performed the analysis on non-spike protein-derived epitopes and compared these to the results above. As expected, a smaller fraction of CD8 T cell recognized epitopes in the SARS-CoV-2 spike protein were found to be conserved compared to those in non-spike proteins (median: 95.1%, range: 92.0-96.6%; versus median: 98.9%, range: 98.0-99.2%; Figure S1C-D). All epitopes overlapping with multi-amino acid substitutions, deletions and/or insertions except one were located in the SARS-CoV-2 spike protein (Figure S1F-G). Such multi-amino acid changes are expected to lead to a lower sequence similarity between altered and reference epitopes. In line with this, there was a significantly lower sequence similarity between mutated and reference CD8 T cell recognized epitopes in the spike protein, compared to the single-amino acid mutations in non-spike proteins (in variants Delta, p=0.043; AY.4.2, p=0.047; Omicron, p=0.028; Figure 4A).

**Figure 4:**
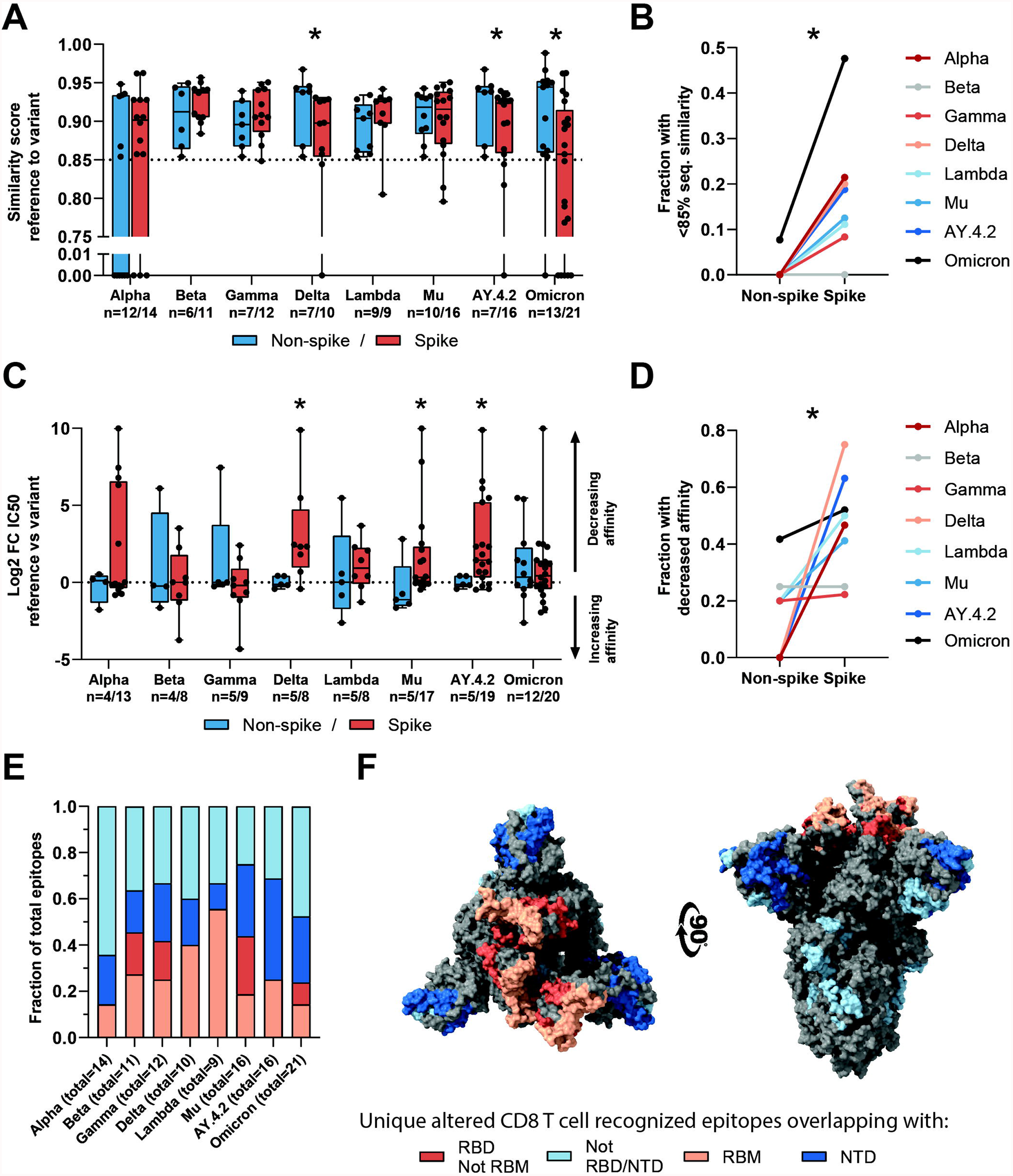
Sequence similarity and predicted binding affinity of spike-versus non-spike-derived mutated CD8 T cell recognized epitopes. **(A)** Box plot comparing the sequence similarity of the altered spike and non-spike protein-derived CD8 T cell recognized epitopes, to the reference epitopes. Sequence similarity of indicated epitopes is set to zero as a result of one or more deletions/insertions in the epitope sequence (Alpha, n = 6/3; Delta, n = 0/2; AY.4.2, n = 0/2; Omicron, n = 1/12). **(B)** Fraction of spike versus non-spike protein-derived epitopes where the sequence similarity of the epitope to the reference epitope dropped below 85% as a result of the mutation. **(C)** Box plot comparing the log2 fold change in predicting binding affinity of spike and non-spike protein-derived CD8 T cell recognized epitopes to the corresponding HLA restriction element, as a result of the mutation. **(D)** Fraction of spike versus non-spike protein-derived CD8 T cell recognized epitopes where the predicted binding affinity of the epitope to the corresponding HLA restriction element was decreased (≥2-fold change in IC50) as a result of the mutation. **(E)** Fractions of unique spike protein-derived CD8 T cell recognized epitopes overlapping with a mutation located in the N-terminal domain (NTD), receptor-binding domain (RBD), receptor-binding motif (RBM) or a mutation located outside of these domains. **(F)** SARS-CoV-2 spike trimer in the open conformation with one erect RBD. Colors represent unique altered CD8 T cell recognized epitopes overlapping with the indicated domains. Image produced with ChimeraX using PDB accession: 6ZGG. Box plots indicate the median (line), 25th and 75th percentile (box), min and max (whiskers), and all data points (single circles). Statistical significance was tested with a two-tailed Mann–Whitney *U* test (A, C) or with a two-tailed Wilcoxon matched-pairs signed rank test (B, D). Variation in numbers of epitopes in the analyses are due to inclusion of epitopes binding one or more HLA restriction elements. * P < 0.05. n indicates the number of epitopes analyzed per group.

Across the investigated SARS-CoV-2 variants, the fraction of CD8 T cell recognized epitopes with low (<85%) sequence similarity to reference epitopes was significantly higher in spike versus non-spike protein-derived epitopes (p=0.016, Figure 4B). Additionally, mutations in the spike protein of the Delta, Mu and AY.4.2 variants were more detrimental to predicted HLA binding affinity compared to non-spike protein mutations (Delta: p=0.019, Mu: p=0.025, AY.4.2: p=0.030; Figure 4C). In accordance with this, the fraction of CD8 T cell recognized epitopes with decreased predicted binding affinity was significantly higher in spike versus non-spike protein-derived epitopes across the variants (p=0.016, Figure 4D). However, despite these statistically significant differences, these results are derived from a limited number of unique mutated CD8 T cell recognized epitopes per variant.

The overrepresentation of altered CD8 T cell recognized epitopes with multiple amino acid changes, insertions and deletions in the spike protein is clearly pronounced. Accordingly, these epitopes are more profoundly affected in terms of epitope sequence similarity, predicted binding affinity and immunogenicity compared to non-spike protein derived epitopes. These results may be unrelated to T cell immunity and may be explained for example by the high rate of evolution of the spike protein due to natural selection pressure by antibody responses. In line with this, a substantial part (median: 65%, range: 36-75%) of the unique spike protein-derived CD8 T cell recognized epitopes that overlap mutations were located in key domains that are associated with cell attachment and are antibody targets (N-terminal domain, NTD; receptor-binding domain, RBD; receptor-binding motif, RBM; Figure 4E). Moreover, they are primarily present on the surface of the spike protein, making them accessible to host antibodies (Figure 4F).

## 4 Discussion

After the initial SARS-CoV-2 outbreak, SARS-CoV-2 variants Alpha, Delta and Omicron have replaced the previous variant as the globally dominant SARS-CoV-2 variant (31,32). This is the result of accumulated mutations resulting in amino acid changes that have allowed these variants to evade immunity in the general population. This notion is supported by for example data showing that serum from vaccine-recipients is less effective at neutralizing SARS-CoV-2 variants Delta and Omicron (23,24). The mutations do not appear to prevent general cross-recognition of SARS-CoV-2 variants by T cells induced by reference SARS-CoV-2 (25–29). However, systematic data regarding the effect of SARS-CoV-2 variant-specific amino acid changes on the properties of the previously recognized CD8 T cell epitopes has been lacking.

Our analyses revealed that the vast majority of both the spike (median: 95.1%, range: 92.0-96.6%) and pan-proteome (median: 98.9%, range: 98.0-99.2%) derived CD8 T cell recognized epitopes were conserved in the investigated SARS-CoV-2 variants. In accordance with the experimental data described above, this suggests that memory T cell responses are not likely to be diminished upon re-infection by a different SARS-CoV-2 variant, or upon infection by one of the SARS-CoV-2 variants after vaccination. In addition, for the minority of presented CD8 T cell recognized epitopes that is altered by mutations, the high degree of sequence similarity to the reference epitopes will likely also not prevent cross-recognition by memory CD8 T cells.

The finding that CD8 T cell epitopes from SARS-CoV-2 were generally conserved is in accordance with the concept of antigenic drift. Antigenic drift driven by selective pressure from T cells is largely irrelevant in acute viral infections such as SARS-CoV-2 due to the huge polymorphism of HLA loci in a population and the diverse antigen repertoire these complexes present to T cells (9). Antigenic drift is likely to have a stochastic influence on T cell epitopes -a ‘bystander effect’. Our observations are in line with this notion. First, for the minority of CD8 T cell recognized epitopes that overlap with mutations, epitope sequences are generally conserved in terms of sequence similarity to the reference sequence. Second, the majority (median: 55.6%, range: 46.2-78.6%) of these CD8 T cell recognized epitopes are predicted to possess unchanged or even increased binding affinity to the HLA allele as a result of the mutation. Third, the mutations in the CD8 T cell recognized epitopes in SARS-CoV-2 variants AY.4.2 and Omicron are predicted to result in more immunogenic, rather than less immunogenic T cell epitopes. Fourth, many of the observed changes in predicted binding affinity and sequence similarity of mutated CD8 T cell recognized epitopes in comparison to the reference epitopes, are indeed driven by mutations in the spike protein. Finally, the majority (median: 65%, range: 36-75%) of spike protein-derived CD8 T cell recognized epitopes that overlap with a mutation are located in key domains that are frequently recognized by antibody responses and/or are involved in cell attachment (10). Therefore, on the basis of our analysis and as expected, there is no indication of T cell based selective pressure on SARS-CoV-2 leading to alteration of the CD8 T cell recognized epitopes.

As SARS-CoV-2 derived T cell epitopes are not subject to substantial antigenic drift, T cells are likely to remain consistent in their recognition of infected cells. However, the stochastic influence by the mutations focused on the spike protein affects the ability of spike protein-derived T cell epitopes to be presented to the immune system or to be recognized by previously induced T cell responses. This is most pronounced for the SARS-CoV-2 variants Delta, AY.4.2 and Omicron, which are also most efficient at escaping humoral immunity as a result of numerous mutations in the spike protein. These variants also have the largest, albeit overall minor, negative effect on epitope presentation relative to the other SARS-CoV-2 variants. By only targeting the spike protein, the vaccine induced immunity is limited to SARS-CoV-2 T cell epitopes which are most prone to a ‘bystander’ effect as a result of the high mutation rate of the spike protein. Even though the currently approved vaccines only include the spike protein, our data suggest that T cell immunity can protect against severe COVID-19. However, it does seem like a logical approach to develop next generation vaccines incorporating other parts of the SARS-CoV-2 proteome which can lead to a broader T cell response providing protection should the spike protein undergo further changes over time.

## 6 Author Contributions

OI and SK performed the experiments, analyzed the data and wrote the manuscript. PK obtained the funding and supervised the study. All authors contributed to the article and approved the submitted version.

## 8 Data Availability Statement

The datasets analyzed for this study are included in the article/supplementary data. Further inquiries can be directed to the corresponding author.

