## Supplementary Material for "*In silico* analysis predicts a limited impact of SARS-CoV-2 variants on CD8 T cell recognition"

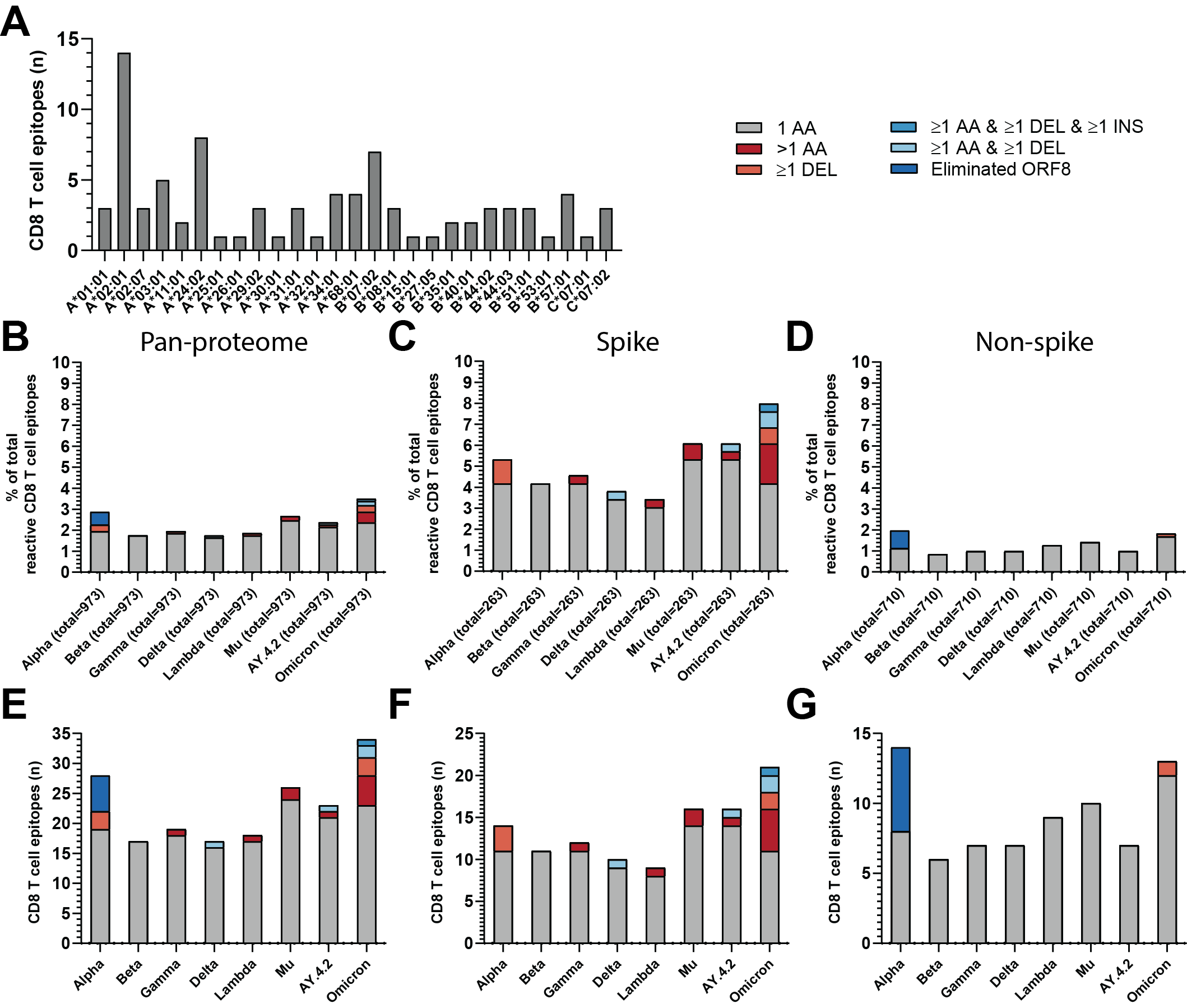

**Supplementary Figure 1.** **CD8 T cell recognized epitopes that overlap with a mutation**

**(A)** Numbers of unique CD8 T cell recognized epitopes included in this study that bind the indicated HLA restriction elements, based on literature. **(B)** Percentage of pan-proteome, spike **(C)**, non-spike **(D)** CD8 T cell recognized epitopes per variant that harbor the indicated types of mutations. **(E)** Numbers of pan-proteome, **(F)** spike, **(G)** non-spike CD8 T cell recognized epitopes per variant that harbor the indicated categories of mutations.

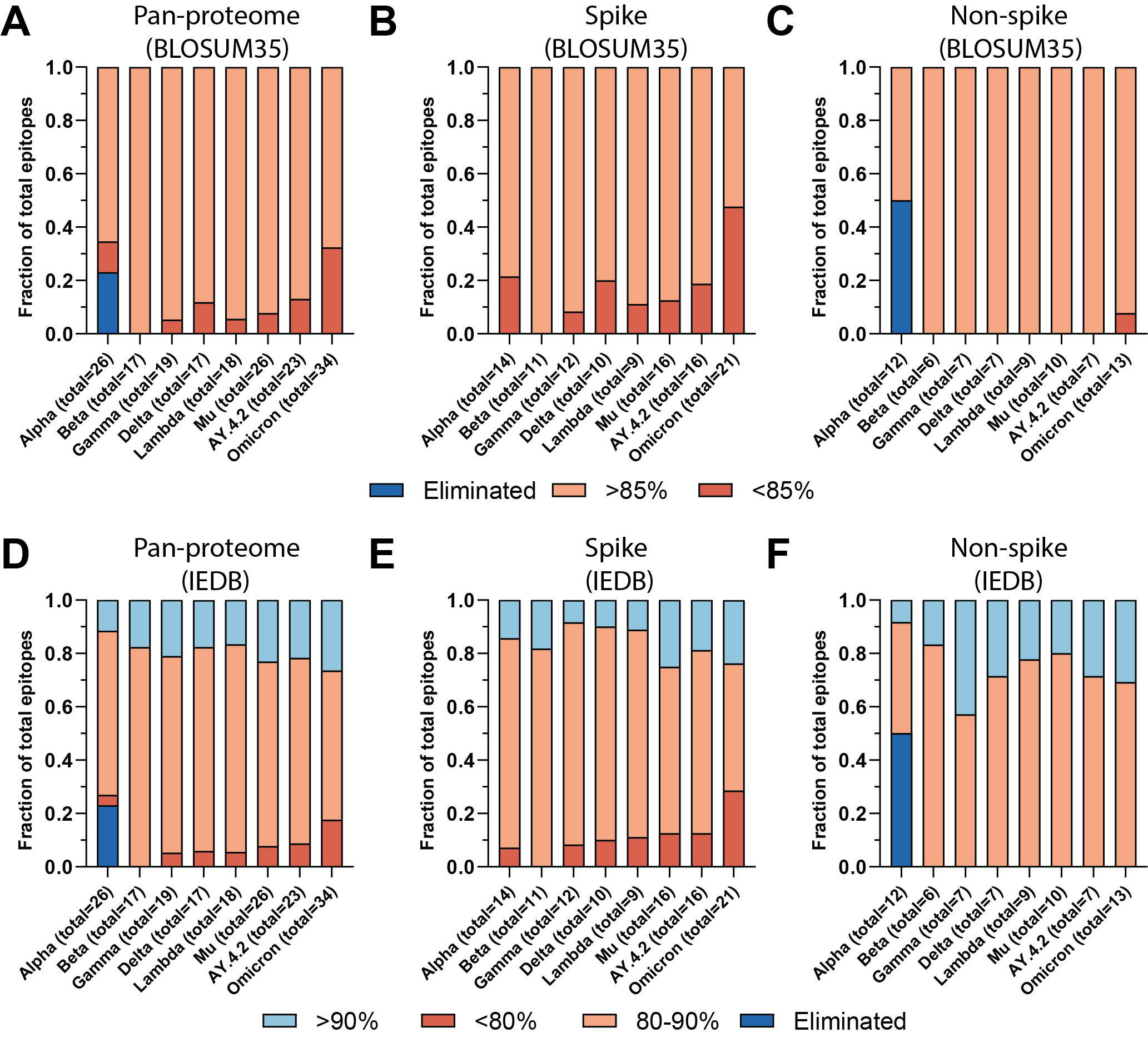

**Supplementary Figure 2.** **Mutant epitope sequence similarity to reference epitope sequence.**

**(A)** Fractions of pan-proteome, spike **(B)**, non-spike **(C)** CD8 T cell recognized epitopes where the sequence similarity of the altered epitope to the reference epitope was below or above 85%, using the method by Frankild et al. described in the text. **(D)** Fractions of pan-proteome, spike **(E)**, non-spike **(F)** CD8 T cell recognized epitopes where the sequence similarity of the altered epitope to the reference epitope was below 80%, above 90% or between 80-90%, using the IEDB epitope clustering tool described in the text. Epitopes were considered eliminated as a result of the ORF8 Q27* stop codon mutation (Alpha, n = 6).

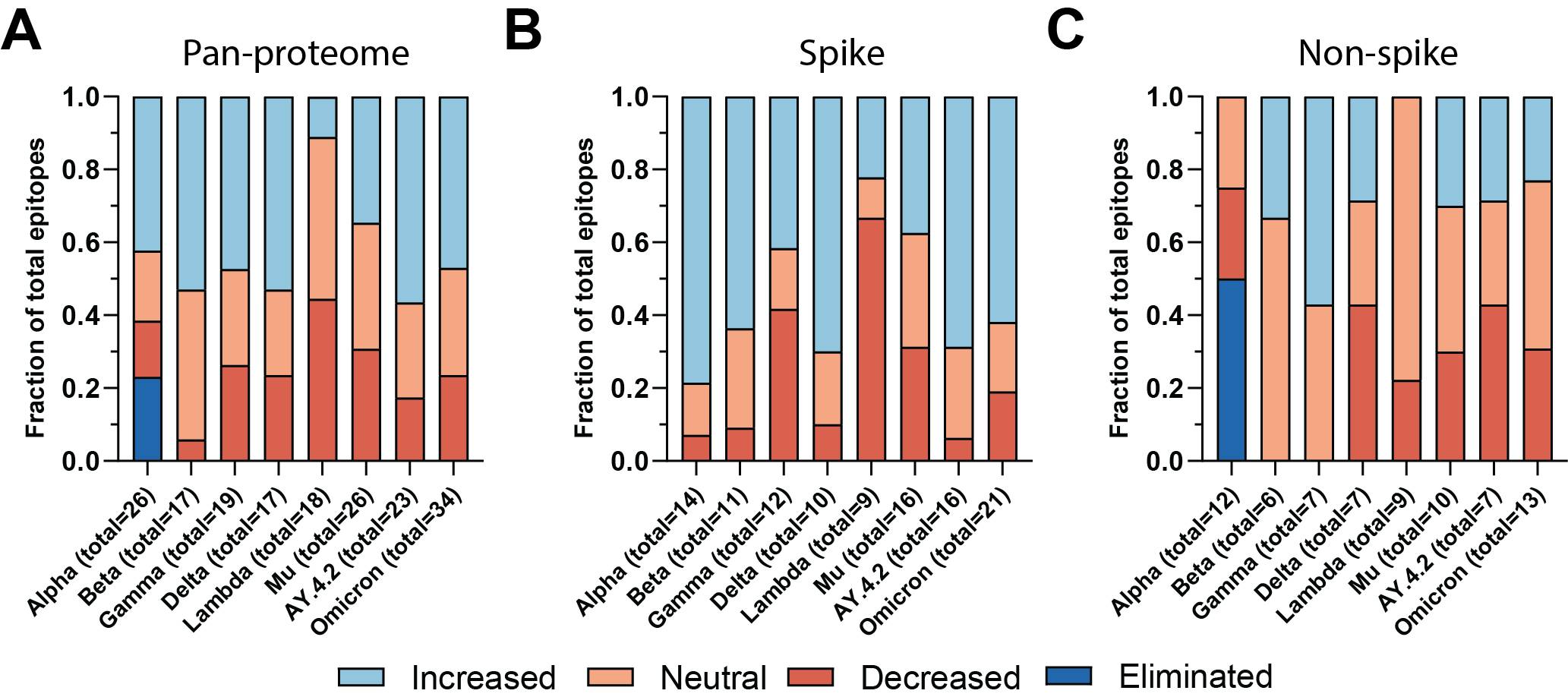

**Supplementary Figure 3.** **Fraction of altered CD8 T cell recognized epitopes with a change in predicted immunogenicity**

**(A)** Fractions of pan-proteome, spike **(B)**, non-spike **(C)** CD8 T cell recognized epitopes where the predicted immunogenicity increased, decreased or was unchanged as a result of the overlapping mutation.

|  | **SARS-CoV-2 variant** | | | | | | | |
| --- | --- | --- | --- | --- | --- | --- | --- | --- |
|  | **Alpha** | **Beta** | **Gamma** | **Delta** | **Lambda** | **Mu** | **AY.4.2** | **Omicron** |
| **ORF1ab** | T1001I, ***A1708D***, ***I2230T***, DEL3675/3677, ***P4715L*** | T265I, ***K1655N***, K3353R, DEL3675/3677, ***P4715L*** | ***S1188L***, K1795Q, DEL3675/3677, ***P4715L***, E5665D | A1306S, P2046L, P2287S, ***V2930L***, T3255I, ***T3646A***, ***P4715L***, G5063S, P5401L, A6319V | T1246I, P2287S, ***F2387V***, ***L3201P***, T3255I, G3278S, DEL3675/3677, ***P4715L*** | ***T1055A***, ***T1538I***, T3255I, Q3729R, ***P4715L***, P5743S | A1306S, P2046L, P2287S, A2529V, ***V2930L***, T3255I, ***T3646A***, ***P4715L***, G5063S, P5401L, A6319V | ***K856R***, S2083I, DEL2084/2084, ***A2710T***, T3255I, P3395H, ***DEL3674/3676***, I3758V, ***P4715L***, I5967V |
| **S** | ***DEL69/70***, ***DEL144/144***, ***N501Y***, A570D, ***D614G***, ***P681H***, ***T716I***, ***S982A***, D1118H | ***D80A***, ***D215G***, ***K417N***, ***E484K***, ***N501Y***, ***D614G***, ***A701V*** | ***L18F***, ***T20N***, ***P26S***, ***D138Y***, R190S, ***K417T***, ***E484K***, ***N501Y***, ***D614G***, ***H655Y***, ***T1027I***, ***V1176F*** | ***T19R***, ***E156G***, ***DEL157/158***, ***L452R***, T478K, ***D614G***, ***P681R***, D950N | ***G75V***, ***T76I***, ***R246N***, ***DEL247/253***, ***L452Q***, ***F490S***, ***D614G***, ***T859N*** | ***T95I***, ***Y144S***, ***Y145N***, ***R346K***, ***E484K***, ***N501Y***, ***D614G***, ***P681H***, D950N | ***T19R***, ***T95I***, ***G142D***, ***Y145H***, ***E156G***, ***DEL157/158***, ***A222V***, ***L452R***, T478K, ***D614G***, ***P681R***, D950N | A67V, ***DEL69/70***, ***T95I***, ***G142D***, ***DEL143/145***, ***N211I***, ***DEL212/212***, ***INS214EPE***, ***G339D***, ***S371L***, ***S373P***, ***S375F***, S477N, T478K, ***Q493R***, ***G496S***, ***Q498R***, ***N501Y***, ***Y505H***, T547K, ***D614G***, ***H655Y***, ***N679K***, ***P681H***, ***D796Y***, N856K, Q954H, N969K, ***L981F*** |
| **ORF3a** |  | ***Q57H***, S171L | S253P | S26L |  | ***Q57H***, DEL256/257 | S26L |  |
| **E** |  | P71L |  |  |  |  |  | T9I |
| **M** |  |  |  | I82T |  |  | I82T | D3G, ***Q19E***, ***A63T*** |
| **ORF7a** |  |  |  | ***V82A***, T120I |  |  | ***V82A***, T120I |  |
| **ORF7b** |  |  |  | T40I |  |  | T40I |  |
| **ORF8** | ***Q27****, R52I, ***Y73C***, S84L | S84L | S84L, E92K | S84L, DEL119/120 | S84L | ***T11K***, ***P38S***, S67F, S84L | S84L, DEL119/120 | S84L |
| **N** | ***D3L***, R203K, G204R, S235F | T205I | ***P80R***, R203K, G204R | D63G, R203M, G215C, D377Y | ***P13L***, R203K, G204R, G214C | T205I | D63G, R203M, G215C, D377Y | ***P13L***, DEL31/33, R203K, G204R |

**Supplementary Table 1.** **SARS-CoV-2 variant amino acid changes**

***Bold, italic:*** *mutations overlapping with CD8 T cell epitopes*

** stop codon*
